# Replicability of unsupervised deep learning derived image phenotypes

**DOI:** 10.64898/2026.05.19.726257

**Authors:** Tian Xia, Sheikh Muhammad Saiful Islam, Ziqian Xie, Xingzhong Zhao, Degui Zhi

## Abstract

Unsupervised deep-learning image phenotypes derived from brain MRI are propelling imaging genetics to link brain structure to genetic variation. However, their replicability across data sets has not been sufficiently evaluated, raising questions about whether they capture robust biological structure or reflect training-specific artifacts. Here, we assess the replicability of unsupervised deep-learning image phenotypes under variation in model initialization, data partitioning, and cohort, directly evaluating their stability across experimental conditions.

We trained multiple models under (i) different training batch random seeds, (ii) cross-validation splits, and (iii) independent datasets (UKB and ADNI), across CNN and ViT architectures,. We then derived representations from a separate UKB discovery cohort (N = 22,985) for both trained models and random initialized models without training. The representation stability was assessed using centered kernel alignment (CKA; mean ViT 0.74 vs random 0.27) and kernel canonical correlation analysis (KCCA; mean ViT 0.84 vs random 0.60), as well as genetic discovery stability using loci overlap ratio (mean ViT 0.45 vs random 0.08). We further applied weighted MAXVAR generalized CCA to 12 embeddings to extract a shared 30-dimensional subspace. Our result showed that UDIPs exhibit statistically significant stability (CKA, KCCA t test p < 0.001) across training perturbations and preserve biologically meaningful structure (loci overlap ratio t test p <0.001) across cohorts, supporting their use in imaging genetics.

## Introduction

Linking genetic variations to human brain structure is a critical step toward understanding the biological basis of cognition and disease. Progress in this area, however, has been constrained by a key limitation: imaging features are often predefined, which restricts the discovery of novel associations.

In recent work, we developed the framework of unsupervised deep learning derived image phenotypes [1], that can derive informative brain imaging features from a large collection of images beyond the traditional IDPs (imaging-derived phenotypes) that were predefined by human experts. We applied deep learning models, such as convolutional neural networks (CNNs) [1, 2] and Vision Transformer (ViT) [3]-based autoencoders, to derive 128-dimensional representations from brain MRI scans of UK Biobank (UKB) participants. After training in a subset of UKB (n=6,130), these models were subsequently used in an independent discovery subset of UKB (n=22,867) to derive unsupervised deep-learning image phenotypes and for conducting genome-wide association studies (GWAS). We showed that these unsupervised deep-learning image phenotypes were informatively correlated with demographic features, brain structure and diseases, and their GWAS can identify significant genetic variants, many are replicated in another UKB replication subset (n=11,000). This framework has been successfully applied to multimodal of MRI including T1, T2, and DTI (FA) images [1-3].

A major concern regarding unsupervised deep-learning image phenotypes is their reproducibility and replicability. For reproducibility, representations learned by neural networks can vary due to factors such as random weight initialization, data sampling, stochastic optimization, distributed training, quantization errors, hardware type and more. Some of these factors, such as initialization, can be controlled, whereas others are inherently difficult to manage. In particular, stochastic gradient descent might find different local minima or early stopped solutions in parameter space, resulting in different models. In addition, these models were primarily trained and tested only among subsets of UKB, mainly due to limited availability of large brain imaging data sets with genetic data. Even though these subsets are disjointed, they are acquired under unified UKB protocol and undergone unified preprocessing [4]. These limitations point to an important possibility: the learned embeddings capture a combination of biological structure, dataset-specific artifacts as well as stochasticity, which reflect aleatoric uncertainty, epistemic uncertainty and other training or model induced variations [5].

Guided by the framework proposed by the National Academies of Sciences, reproducibility refers to the ability to obtain the same results using the original data and code, whereas replicability refers to the ability to obtain consistent findings when a study is independently repeated using newly collected data or under different experimental conditions, while addressing the same scientific question [6]. This distinction is particularly relevant for deep learning models, where high reproducibility does not necessarily imply strong replicability. For example, Nong et al. conducted an exhaustive literature review on a niche deep learning topic, vulnerability detection, prior to 2022. They reported that although a large proportion of studies (87.5%) were reproducible, only 28.6% of those models remained replicable when retrained and evaluated on a different real-world dataset [7].

In the context of deep learning derived brain MRI traits, only a limited number of studies have examined replicability, robustness, or generalization of deep learning–based representations, and these have largely been conducted in supervised settings. In this literature, robustness is distinguished from replicability by whether the model is retrained: robustness typically refers to training a model on one dataset and directly evaluating it on an external dataset without further retraining, whereas replicability involves retraining the model under new data or conditions. For instance, AlzCLIP [8], trains an Alzheimer’s disease prediction model using both Alzheimer’s Disease Neuroimaging Initiative (ADNI) images and sequencing data and evaluates it on ADNI and UKB. While it demonstrates comparable prediction performance across datasets (AUC = 0.80 for ADNI and 0.81 for UKB) and identifies shared significant loci, it does not directly assess the similarity or stability of the learned embeddings.

Another example is TransferGWAS [9], which evaluates two modelling strategies: a CNN-based autoencoder trained with a combination of supervised and unsupervised objectives, and a model initialized from an ImageNet-pretrained checkpoint. Both are applied to the UKB dataset, and their performance is compared based on the overlap in downstream genetic discoveries. This setup can be interpreted as a broader assessment of replicability, insofar as the two models represent variations of the same embedding derivation framework. However, the comparison remains confined to supervised or hybrid settings and relies solely on overlap-based metrics, without directly assessing the stability of the learned representations themselves.

Despite these efforts, no prior study has rigorously evaluated the replicability of imaging phenotypes across different data sets derived purely from unsupervised learning. Existing deep learning work either evaluates representations indirectly through downstream tasks (e.g., prediction accuracy [8, 10], identified biomarkers [8, 11]) or relegates replication analyses to supplementary materials, rather than treating them as a central focus. This gap is important, as unsupervised approaches may capture less biased and more intrinsic structure in the data because they do not rely on external labels. However, whether such representations are stable and replicable across datasets remains an open question.

In this study, we explicitly evaluate the replicability of unsupervised deep-learning image phenotypes by systematically varying three key training conditions, which form a comprehensive experimental framework spanning multiple sources of variation that commonly affect representation learning: (i) repeated trainings with different random initialization seeds to assess stochastic variability from python, numpy, PyTorch’s CPU GPU and mini batch, (ii) alternative data splits within the same cohort to quantify sensitivity to sample partitioning, and (iii) training on independent datasets, namely ADNI and UKB, to evaluate cross-cohort generalizability. After training, all representations were derived from a separate held-out subset of UKB, the same UKB discovery cohort (N = 22,985) used in our previously published work[1] which served as the reference evaluation set. As a negative control, we additionally derived unsupervised deep-learning image phenotypes from randomly initialized networks without training on the reference evaluation set, allowing us to establish a baseline of spurious or non-informative structure.

We conducted this analysis across two widely used neural network architectures, CNNs and ViTs, to make conclusions less architecture-specific. To quantify agreement and stability of the learned representations, we employed three complementary metrics: (i) centered kernel alignment (CKA), following Simon Kornblith et al. [12], which captures similarity in pairwise sample relationships and is invariant to orthogonal transformations and isotropic scaling; (ii) canonical correlation analysis (CCA), [13], which measures alignment between subspaces; and (iii) loci overlap ratio, which evaluates consistency at the level of downstream genetic discoveries.

Together, this systematic evaluation across initialization, data partitioning, dataset heterogeneity, and model architecture directly addresses the longstanding question of unsupervised deep-learning image phenotypes’ reproducibility and stability. By integrating representation-level and genetics-level metrics, our framework provides a rigorous assessment of whether these deep-learning image phenotypes constitute a reliable and biologically meaningful resource for imaging genetics analyses, rather than artifacts of specific training conditions.

## Results

### Replicablilty of unsupervised deep-learning image phenotypes measured by the CKA

We first assessed representational similarity among 128-dimensional unsupervised deep-learning image embeddings relative to a regular trained (base) model of UK Biobank cohort using CKA. For each pair of representations *X*and *Y*(matched subjects and feature dimensionality), features were column-wise mean-centered, and linear CKA was computed as

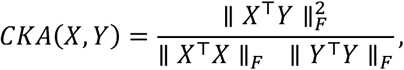

where ∥⋅∥_*F*_ represent Frobenius norm, yielding values in [0, 1], where larger values indicate greater alignment of second-order feature structure. CKA quantifies the agreement between the pairwise similarity structures of two representations. Because it is invariant to orthogonal transformations and isotropic scaling, it provides a robust and comparable measure of representational similarity across models, making it particularly well-suited for evaluating learned embeddings.

For ViT, we used the regular trained base model as the reference and summarized its CKA with four independent training replicates, three cross-validation folds (each relative to fold 0), random-seed control runs (reported as mean ± SD), and ADNI-trained ViT models (mean ± SD across runs). For CNN, the reference was the discovery model. We report CKA with a training replicate, cross-validation fold, ADNI, and random initialization from the same-cohort matrix. ViT replicates and CV folds show high CKA to the base model (≈0.83 and ≈0.81, respectively), ADNI exhibits intermediate similarity (≈0.59), and random controls remain low (≈0.27).

In CNN, similarity to ViT, the pattern is even stronger, with training replicates (≈0.95) and ADNI (≈0.81) exceeding the corresponding ViT values; notably, random CNN baselines (≈0.38) are also higher than those of ViT (≈0.27). This pattern suggests tighter optimization-induced coupling of CNN representations to the reference under CKA, whereas ViT maintains clearer separation between structured runs and random controls.

We tested whether structured conditions exceeded the random-seed baseline. Using per-seed CKA values, we performed two-sided Welch’s *t*-tests comparing the random distribution to each group (replicates, CV folds, ADNI). All comparisons yielded extremely small *p*-values (e.g., random vs. replicates *p* ≈ 2 × 10^−8^, random vs. CV folds *p* ≈ 5 × 10^−5^, random vs. ADNI *p* ≈ 5 × 10^−5^).

### Replicability of unsupervised deep-learning image phenotypes measured by the CCA (kernel)

We assessed alignment between unsupervised deep-learning image phenotypes representations using kernel canonical correlation analysis (KCCA) with a radial basis function (RBF) kernel, implemented via random Fourier features (RFF) to approximate the kernel map at scale (Figure 1). For each pairwise comparison, we summarized agreement by the mean Pearson canonical correlation (ρ) across the first 30 canonical components (Truncation at *K* = 30 was chosen because the first 30 pairs of unsupervised deep-learning image phenotypes replication group has pearson canonical correlation over 0.8)

**Figure 1.**
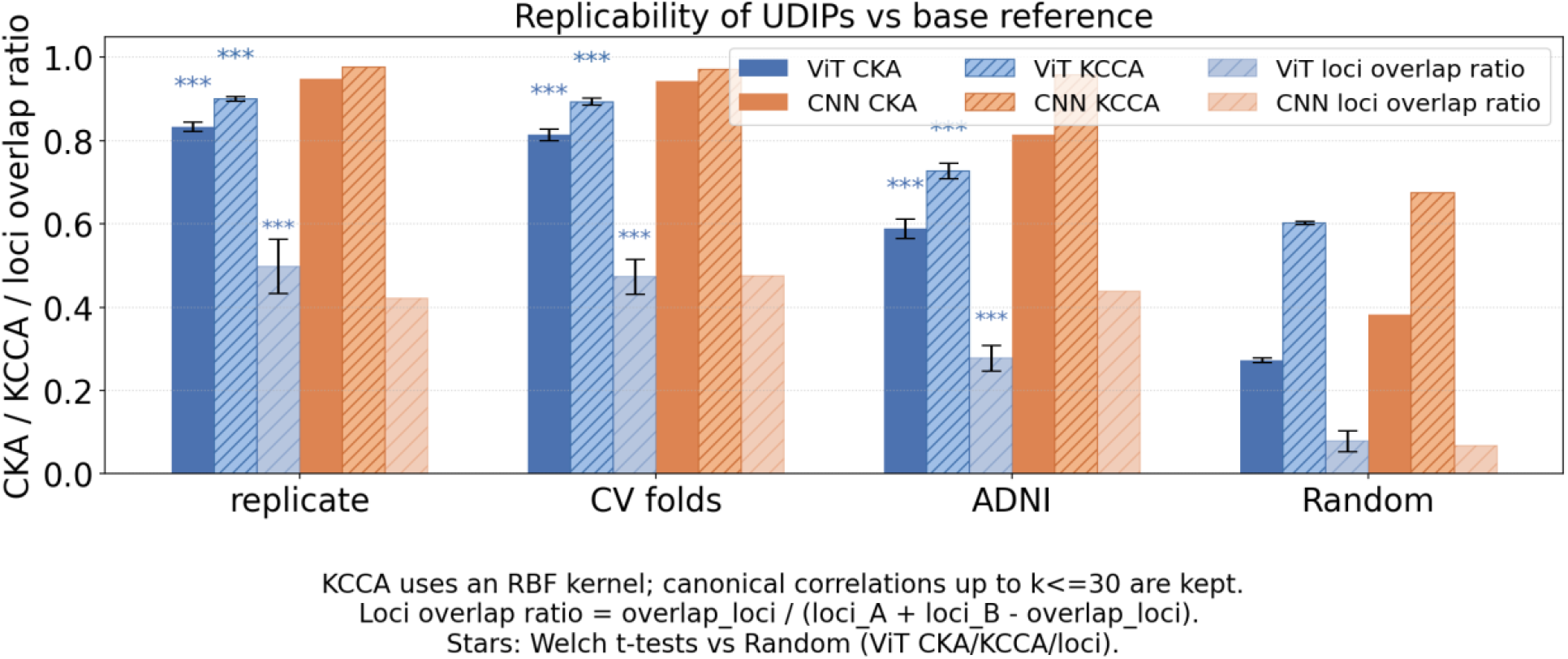
CKA and loci overlap ratio between unsupervised deep-learning image phenotypes derived from models trained with different random seeds, data splits, and distinct datasets for T1 modality.

Within-UKB stability was high when comparing the original UKB embedding to independent UKB replicates trained with different seeds (off-diagonal mean ρ ≈ 0.89–0.91) and when comparing embeddings learned on different cross-validation (CV) folds (off-diagonal mean ρ ≈ 0.88–0.90). In both cases the component-level correlation matrices were nearly uniform, indicating that the leading canonical directions are robust to resampling and to the selection of training subset.

Cross-cohort and negative-control analyses contrasted UKB with ADNI and with matched random inputs. UKB-ADNI agreement remained substantial (mean ρ ≈ 0.73 for the first 30 components), but was materially lower than within-UKB agreement, consistent with cohort-specific acquisition and population differences while still supporting partially shared structure across studies. Relative to these biological comparisons, correlations against random data were markedly reduced (UKB–random and ADNI– random mean ρ ≈ 0.61), indicating that unsupervised deep-learning image phenotypes capture replicable structure that generalizes across cohorts.

Taken together, the first 30 canonical components show strong multivariate alignment between unsupervised deep-learning image phenotypes under stable sampling conditions (ρ often >0.88), supporting the view that, after the appropriate linear transformations induced by KCCA/RFF, many leading components can be brought into close correspondence.

### Replicability of unsupervised deep-learning image phenotypes measured by the loci overlap ratio

We further followed the same genetic analysis procedure as our previous publication of CNN [1] and ViT [3] model to derive genomic loci (see Methods). Briefly, we applied the mixed linear model fastGWA to compute p-values for each unsupervised deep-learning image phenotype, followed by a minP procedure for aggregating multivariate summary statistics and using FUMA to clump significant SNPs (Bonferroni-corrected threshold) into genomic loci. Then each locus was extended by 125 kb and query against each other for determining overlaps. The loci overlap ratio is defined as the overlap loci number divided by the union loci number.

Figure 1 overlays genomic loci overlap ratio on the right axis. For ViT, loci overlap is highest for replicates and CV folds (≈0.50 and ≈0.47), intermediate for ADNI (≈0.28), and low for random (≈0.08), with Welch tests against random loci again showing strong significance (p<0.001). The ViT loci overlap ratio observed between ADNI and UKB is consistent with prior findings based on supervised pretraining and transfer to the UKB dataset for embedding extraction and GWAS analysis [9]. In that study, the overlap ratio between ADNI-supervised pretraining and ImageNet-supervised pretraining was 0.242.

### Find common base of unsupervised deep-learning image phenotypes by Generalized CCA

We further used multi-view maximum-variance (MAXVAR) generalized canonical correlation analysis (GCCA) [14] to find shared space of 12 ViT T1 unsupervised deep-learning image phenotypes embeddings of 128-dimensions. The analysis extracted *K* = 30 shared subject-space directions *G* from 12 views: the UKB discovery model, four independent training replicates, three cross-validation folds, and four ADNI-trained models. All design matrices were column-wise *z*-scored prior to analysis.

Let 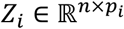 be the matched-subject embedding matrix for view *i*, 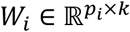 the per-view projection, and *G* ∈ ℝ^*n*×*k*^ the shared latent representation (here *k* = 30 for the common-space analysis). The MAXVAR objective solved is:

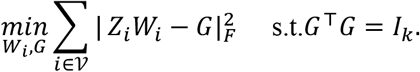

This finds a common subspace where all 12 projected views align as closely as possible to the same *G*, which is then used for explained-variance and correlation diagnostics. To avoid imbalance between cohorts (eight UKB-family views vs. four ADNI-family views), we applied view-level reweighting in the MAXVAR objective so that UKB and ADNI contributed equally overall: each UKB-family view received weight 1/8, and each ADNI view 1/4. This prevents the larger number of UKB runs from dominating the learned shared subspace.

For each view *i*, we report the fraction of Frobenius variance captured by projecting *Z*_*i*_*W*_*i*_ onto the shared subspace:

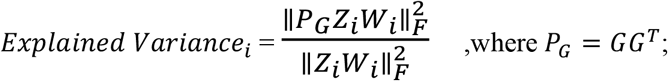

*Z*_*i*_*W*_*i*_ ∈ ℝ^*n*×*d*^ is the view-specific representation in the sample space. *G* ∈ ℝ^*n*×*k*^ spans the shared subspace. *P*_*G*_ = *GG*^⊤^ is projection matrix of *ZW* to *G* in the sample space with the size of *n* by *n*, where *n* is the sample size. ∥⋅∥_*F*_ denotes the Frobenius norm.

Geometrically, this quantity is equivalent to a variance-weighted average of cos^2^of the principal angles between the subspace spanned by *Z*_*i*_*W*_*i*_and *G*. Accordingly, values approaching 1 indicate strong subspace alignment (small principal angles), whereas values near 0 indicate orthogonality.

The results in Figure 2 indicate that all unsupervised deep-learning image phenotypes variants, despite differences in training runs, folds, or cohorts, retain a substantial shared low-dimensional factor structure captured by the 30-dimensional GCCA subspace. This suggests that the learned embeddings are not independent representations, but rather different parameterizations of a common latent manifold that can be aligned through linear projection.

**Figure 2.**
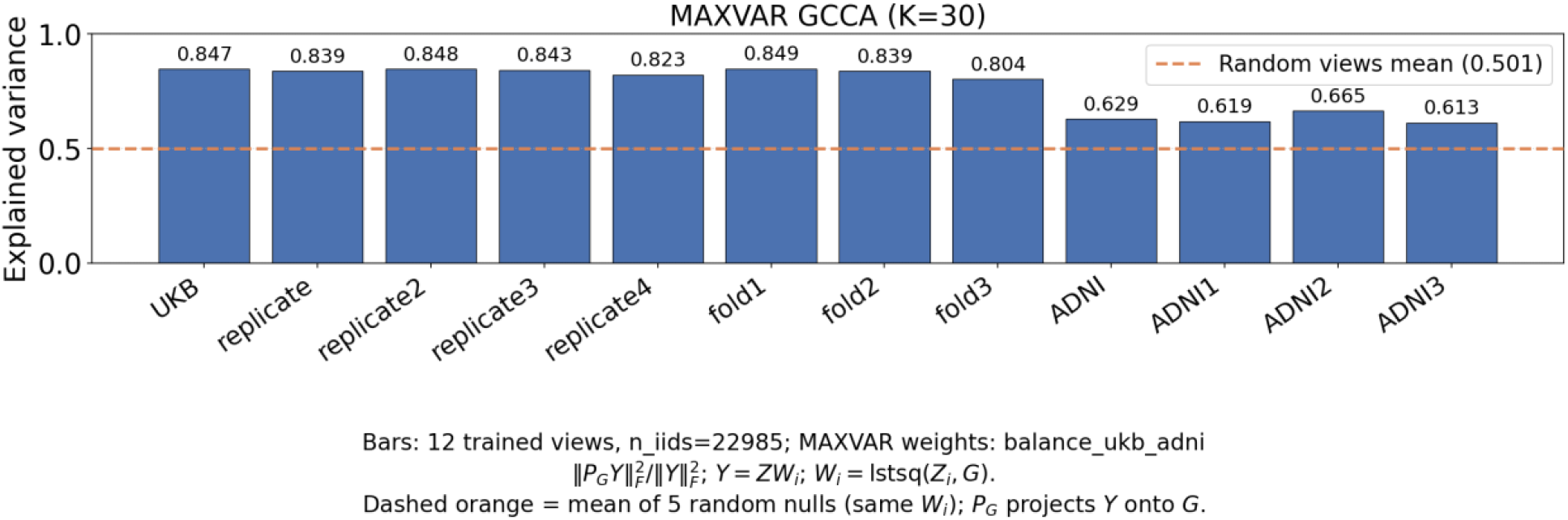
Generalized canonical correlation analysis (GCCA; MAXVAR, *K* = 30, balanced UKB-ADNI weights): explained variance of the predicted embedding *Y*_*i*_ = *Z*_*i*_*W*_*i*_ in the shared 30-D subspace span(*G*) for all groups.

While the degree of alignment varies across cohorts. UKB-based models are more tightly embedded in this shared space than ADNI models, which exhibit larger rotational deviation. All trained models outperform random projections in both overall performance (Figure 2) and breadth of representation (detailed projections for each GCCA dimension are provided in Supplementary Figure 1). In summary, within this framework, a linear mapping exists that relates any two representations through their projections onto the shared subspace, but the residual (off-subspace) components are larger for cross-cohort transfers, reflecting imperfect but still meaningful shared structure across unsupervised deep-learning image phenotypes.

## Discussion

Our results demonstrate strong concordance of unsupervised deep-learning image phenotypes across experimental conditions, supporting their replicability and transferability for broader applications, such as the Alzheimer’s Disease Sequencing Project. Across complementary metrics, we observe consistent evidence that independently trained models converge to a shared representational structure: high CKA and CCA agreement within UKB, and across ADNI, significant separation from random controls, and reproducible GWAS loci with substantial overlap across replicates and cross-validation folds. The weighted MAXVAR GCCA analysis further shows that this agreement is not limited to pairwise similarity but extends to a common low-dimensional subspace that captures comparable variance across models, indicating that unsupervised deep-learning image phenotypes encode stable, biologically meaningful factors rather than idiosyncratic training artifacts. Together, these findings support a view in which unsupervised deep-learning image phenotypes are replicable within-cohort and across cohorts (Table 1), making them suitable for downstream genetic analyses and integrative studies, while also motivating continued attention to cohort-specific variation when deploying these representations in heterogeneous datasets.

**Table 1.** Overview of ViT unsupervised deep-learning image phenotypes replicability.

| metric | ViT<br>(replicate+cvfold+ADNI) | Random |
| --- | --- | --- |
| CKA | $0.7388 \pm 0.1205$ | $0.2729 \pm 0.0062$ |
| KCCA | $0.8353 \pm 0.0868$ | $0.6028 \pm 0.0040$ |
| GCCA | $0.7610 \pm 0.1042$ | $0.4049 \pm 0.1692$ |
| Loci overlap ratio | $0.4466 \pm 0.1014$ | $0.0781 \pm 0.0250$ |

One issue worth clarifying is the distinction among several closely related concepts: reproducibility (or repeatability), replicability, and robustness (or generalizability). Conventionally, repeatability refers to whether the same person can obtain the same result using the same data and tools, whereas reproducibility refers to whether a different person can achieve the same result under the same conditions. These definitions are relatively clear. Replicability is traditionally defined as obtaining consistent results using different data but the same methods, while robustness refers to obtaining consistent results using [15]. However, these definitions become oversimplified in the context of deep learning. In deep learning pipelines, data usage typically involves two stages: training and inference. Importantly, the training data are effectively encoded into the model parameters, blurring the distinction between “data” and “tool.”

As a result, a single deep learning framework can be used to evaluate both replicability and robustness, depending on how the experiment is designed. For instance, training the same model architecture on independent datasets such as ADNI and UKB and then comparing the learned representations constitutes a test of replicability (same method, different data). In contrast, applying models trained on different datasets (i.e., different “tools”) to a common test dataset (e.g., UKB) and comparing the resulting embeddings reflects robustness under the traditional definition.

To clarify these distinctions, we summarize the relevant definitions within a deep learning framework by explicitly separating the training and inference stages (Table 2). Repeatability and reproducibility retain their standard meanings in both stages. During training, using the same model with different datasets is defined as replicability. During inference, applying a fixed pretrained model to new in-distribution or out-of-distribution datasets is characterized as robustness [16].

**Table 2.** Definition of terms on deep learning model inference stage.

| Train/Retrain/Finetune | Same data | Different data |
| --- | --- | --- |
| Same model | Repeatability /<br>Reproducibility | Replicability |
| Different model | Model Consistency | Transferability |
| Inference | Same data | Different data |
| Same model | Repeatability /<br>Reproducibility | Robustness |
| Different model | Model Consistency | Generalizability |

Following this definition, we assessed the model consistency of unsupervised deep-learning image phenotypes across CNN and ViT architectures (Supplementary Figure 2). The CKA, kernel CCA, and loci overlap ratios are higher than those observed in the random baseline (CKA 0.82 vs 0.29; KCCA 0.91 vs 0.61; loci overlap ratio 0.43 vs 0.05) and fall within the range expected from replicability analyses, indicating strong model consistency.

We also explored whether robustness or replicability provides a more reliable evaluation metric by analyzing embedding variance (Supplementary Figure 3). We found that the ADNI-trained model exhibited higher embedding variance when inference was performed on ADNI data, whereas the UKB-trained model showed higher embedding variance when evaluated on UKB data. This suggests that each trained model is somewhat biased toward, and more sensitive to, its own inference dataset. Therefore, robustness based on embedding variance does not appear to be a stable metric. In contrast, when the inference dataset is held constant, the observed trend remains consistent regardless of whether inference is performed on ADNI or UKB data, indicating that replicablity is a better metric.

Regarding the GWAS pipeline, we acknowledge that the min-P procedure is not an optimal multivariate association test. However, it provides a conservative lower bound on the number of detectable genetic associations, as it requires a locus to show strong evidence of association in at least one component after correction for multiple correlated tests. In the literature, while min-P has been widely adopted as a baseline approach for multi-trait GWAS, several more powerful multivariate methods have since been developed to improve locus discovery, including MTAG [17], MOSTest [18]and C-GWAS [19]. In this study, we employed min-P primarily to enable a fair comparison with our previous published CNN-based method, which used the same procedure. Investigation of alternative multivariate GWAS frameworks is an important direction for future work.

Regarding the genetic overlap measurement, complementary metrics on top of loci overlap ratio such as genetic correlation offer a more comprehensive view. Loci overlap may only capture a limited portion of the underlying polygenic architecture, as it is restricted to genome-wide significant hits and therefore reflects only the most detectable associations. Because complex brain traits are highly polygenic, a substantial proportion of the genetic signal is distributed across many variants with small effects that do not reach genome-wide significance in standard GWAS. Consequently, relying solely on loci overlap may underestimate the true degree of shared genetic information across unsupervised deep-learning image phenotypes. However, estimating genetic correlation at this scale is computationally intensive and time-consuming, and will therefore be incorporated in future work.

Another limitation of current study is we only used UKB discovery cohort (N=22,985) as the reference evaluation data set. This is mainly a practical concern as UKB sample size is much greater to offer sufficient power. Future studies should further evaluate these models in other reference evaluation data sets.

Our framework for rigorously evaluating unsupervised models is broadly applicable across domains. It can be extended to other brain MRI modalities, such as T2-FLAIR and FA images, as well as to imaging data from other organs, including the heart [20]and the retina [21]. Beyond imaging, the framework may also generalize to non-imaging modalities such as EEG waveforms [22]. In addition, it is not limited to unsupervised settings and can be applied to supervised models [8], as well as self-supervised approaches such as contrastive learning [23].

## Methods

### UK Biobank

The UK Biobank [24] is a nationwide population resource encompassing approximately 500,000 participants recruited across the United Kingdom, with phenotypic, imaging, and genetic data collected between 2006 and 2010. The study operates under established ethical approval, with governance and oversight details available at https://www.ukbiobank.ac.uk/learn-more-about-uk-biobank/governance/ethics-advisory-committee. In total, 29,115 brain MRI scans were included in this study. Model development used 4,597 scans for training and 1,533 for validation. Learned representations were subsequently extracted from the remaining 22,985 individuals, corresponding to the same “discovery cohort” used in prior brain imaging GWAS analyses [25, 26].

For genetic analyses, imputed genotype data from the UK Biobank [24] underwent standard quality control procedures. Variants were retained if they met thresholds of minor allele frequency > 0.0001, resulting in 8,925,988 single-nucleotide polymorphisms in the discovery cohort.

### Alzheimer’s Disease Neuroimaging Initiative (ADNI) [27]

Data used in the preparation of this article were obtained from the Alzheimer’s Disease Neuroimaging Initiative (ADNI; including ADNI-1, ADNI-2, ADNI-3, and ADNI-GO; downloaded in February 2025) database (adni.loni.usc.edu). The ADNI was launched in 2003 as a public-private partnership, led by Principal Investigator Michael W. Weiner, MD. The primary goal of ADNI has been to test whether serial magnetic resonance imaging (MRI), positron emission tomography (PET), other biological markers, and clinical and neuropsychological assessment can be combined to measure the progression of mild cognitive impairment (MCI) and early Alzheimer’s disease (AD). A total of 2,594 MRI scans from 1,524 subjects were included in this study.

### Model

#### Model Input Datasets and Preprocessing

UK Biobank MRIs were downloaded on October 15, 2021. UK Biobank participants had birth year between 1934 to 1971 with female to male ratio of 0.52. UK Biobank has provided a bias-field-corrected version of the brain-extracted T1-weighted (T1) captured mainly using standard Siemens Skyra 3T running VD13A SP4 (as of October 2015), with a standard Siemens 32-channel RF receive head coil. Resolution of T1 is 1 × 1 × 1 mm (https://biobank.ctsu.ox.ac.uk/crystal/crystal/docs/brain_mri.pdf).

To promote generalizability and minimize manual feature engineering, we adopted the standard preprocessing pipeline developed by the UK Biobank MRI team. Brain MRI preprocessing was performed primarily using FSL (https://www.fmrib.ox.ac.uk/ukbiobank/), and included defacing, brain extraction with BET, linear and non-linear registration to standard space using FLIRT and FNIRT, and bias-field correction using FAST. Bias-field-corrected, brain-extracted T1-weighted images were retained for analysis. All images were subsequently linearly registered to MNI152 space using affine transformation with 12 degrees of freedom via UK Biobank-provided precomputed matrices in FLIRT. Linear registration was chosen to normalize head size and achieve cross-subject alignment while preserving subject-specific structural deformations, in contrast to non-linear registration. The resulting linearly registered, defaced, bias-field-corrected images were used in all downstream analyses. Image intensities were normalized on a per-scan basis using Z-score normalization. Following affine registration, T1 weighted volumes had dimensions of 182 × 218 × 182 voxels and were zero-padded to 182 × 224 × 182 to enable partitioning into equal-sized patches.

To construct a high-quality deep learning dataset, we leveraged UK Biobank’s precomputed MRI quality metrics: the inverted contrast-to-noise ratio (Data Field 25735) and the discrepancy between T2 FLAIR and T1 brain images (Data Field 25736). Images with values below the 95th percentile for both metrics-corresponding to higher-quality scans-were retained. For participants with multiple visits, only the first scan was kept to ensure consistency across the dataset. This filtering resulted in a cohort of 6,130 scans from individuals of diverse ethnic backgrounds. The dataset was randomly split into a training set of 4,597 images (75%) and a validation set of 1,533 images (25%). The validation set was reserved exclusively for hyperparameter tuning and model checkpointing during training.

For the ADNI data, we downloaded 2,594 T1-weighted structural MRI scans acquired on 3T scanners in NIfTI format, each with vendor-provided on-scanner intensity non-uniformity correction. All images were processed using an in-house preprocessing pipeline to ensure consistency across sites and acquisition protocols. First, skull stripping was performed using DeepBET (threshold = 0.5) to remove non-brain tissue while preserving cortical boundaries [28]. Next, images were aligned to MNI152 template space using affine registration implemented in ANTs [29], enabling cross-subject anatomical alignment. Finally, N4 bias field correction (implemented in ANTs) was applied to further reduce residual intensity inhomogeneity. This standardized preprocessing pipeline facilitates robust downstream feature extraction and cross-cohort comparability. The final T1 weighted volumes had dimensions of 182 × 218 × 182 voxels and resolution of 1 × 1 × 1 mm and were zero-padded to 182 × 224 × 182 to enable partitioning into equal-sized patches. The dataset was randomly split into a training set of 1,945 images (75%) and a validation set of 649 images (25%).

#### Model training

We adopt CNN [1]and ViT [3] autoencoder framework from our previous publication. Details could be found in our previous paper.

Briefly, The CNN architecture is an autoencoder with CNN block of 3 × 3 × 3 kernel and 2 × 2 × 2 kernel for max pooling. The model consist of 5 encoding CNN blocks, an average pooling layer with dimension 128 and 5 decoding CNN blocks. The model is trained for 100 epochs and converge around 20 epochs.

The ViT-AE architecture consisted of encoders, an average pooling layer and decoders. The raw images are partitioned into patches and embedded into 128 dimensions, fed into the vision transfomer along with its position embeddings. The combined token embeddings passed through 12 encoding transformer layers, the average pooling layer, 12 decoding transfomrer layers and were finally decoded by a linear projection to reconstruct the input patches. MSE loss was computed between predicted patches and original patches within the batch mask. The learning schedule is a CosineAnnealingWarmRestarts learning rate scheduler with T_0=10, T_mult=2, eta_min=1×10-6 and AdamW optimizer with an initial learning rate of 0.001. The ViT-AE model was trained for 300 epochs on the training set and validated on a separate validation set on 4 Nvidia H100 GPUs.

#### Model embedding extraction

We froze the CNN/ViT encoder and average pooling layer and remove the decoder layer, then applied the model to T1 weighted MRIs of 22,985 held-out individuals (discovery cohort) to get the representations of size 22,985 samples × 128 dimensions from the average pooling layer.

### Genetic analyses

We implemented a comprehensive quality-control pipeline on the downloaded genetic data, following our previous work [1], resulting in 8,469,833 single-nucleotide polymorphisms (SNPs) across 22,985 individuals of white British ancestry. Briefly, duplicated variants across the 22 autosomal chromosomes were removed, and individuals whose genetically inferred sex did not match their self-reported sex were excluded. Additional filtering criteria removed variants with a minor allele frequency (MAF) below 0.01% or a genotyping missing rate exceeding 5%.

#### GWAS

GWAS were conducted on 22,867 individuals for 128 extracted unsupervised deep-learning image phenotypes. Associations between SNPs and unsupervised deep-learning image phenotypes were tested using linear mixed models implemented in FastGWA [30] within GCTA (v1.94.1), with a MAF threshold of 0.01. Covariates included age (Field ID: 21003), age^2^, sex (Field ID: 31), sex × age, sex × age^2^, the first 10 genetic principal components (Field ID: 22009), intracranial volume (Field ID: 25000), inverted contrast-to-noise ratio (Field ID: 25735), head positioning in the scanner (Field IDs: 25756-25758), scanner table position (Field ID: 25759), assessment center location (Field ID: 54), and date of assessment (Field ID: 53). For participants with multiple visits, only the first scan was retained. A Bonferroni-corrected significance threshold of 5 × 10^−8^/128 was applied, and summary statistics were compiled with the most significant p-value (minP) reported for each SNP.

#### Annotation of genomic loci

Genomic loci were annotated using FUMA [31]. Lead SNPs were first identified based on linkage disequilibrium (r^2^ ≤ 0.1) and physical proximity (<250 kb), and assigned to non-overlapping genomic loci. Within each locus, the SNP with the lowest p-value (the top lead SNP) was used to represent the locus.

#### Overlap of genomic loci

To assess locus overlap, each identified locus was extended by 125 kb on either side, and interval trees were constructed for each chromosome. Loci from our analysis were then queried against these interval trees to determine overlapping regions.

## Data Availability

The previous published CNN and ViT model codes as well as checkpoints are publicly accessible at Github at https://github.com/ZhiGroup/DeepENDO, https://github.com/ZhiGroup/UDIP-ViT. The comparison experiments conducted in this manuscript are publicly accessible at https://github.com/ZhiGroup/UDIP-Replicability.

The unsupervised deep-learning image phenotypes and GWAS summary statistics are available upon reasonable request after removing sensitive UKbiobank patient information. Genomic loci annotation used data from FUMA (https://fuma.ctglab.nl/). Individual data from UKBB can be requested with proper registration at https://www.ukbiobank.ac.uk/. All unrestricted data supporting the findings are also available from the corresponding author upon request.

## Acknowledgements

This work was supported by grants from the National Institute on Aging U01AG070112 and R01AG081398.

We used a large language model to assist with language editing and clarity of presentation. The authors reviewed and edited all content and take full responsibility for the accuracy and integrity of the work.

## Ethics declarations

### Ethical Approval

Our analysis was approved by the UTHealth Houston committee for the protection of human subjects under No. HSC-SBMI-20-1323. UKBB has secured informed consent from the participants in the use of their data for approved research projects. UKBB data was accessed via approved project 24247.

