## Supplementary Figures for "Replicability of unsupervised deep learning derived image phenotypes"

**Supplemental Figures for Replicability of unsupervised learning derived image phenotypes (UDIPs)**

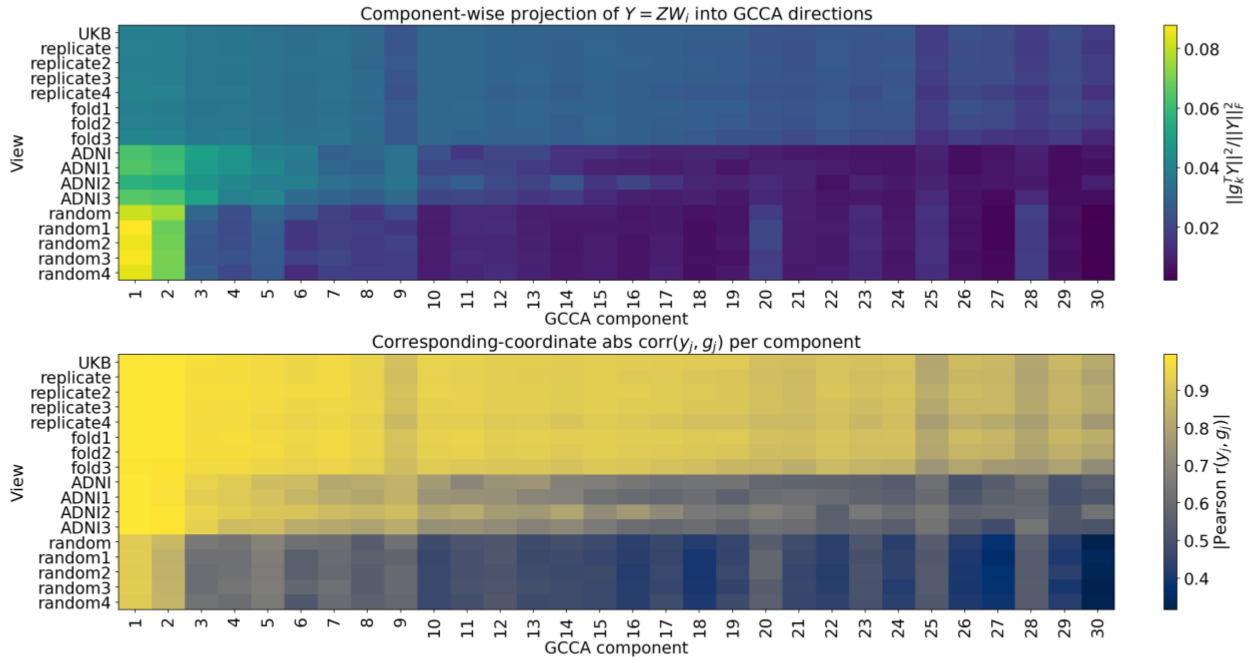


**Supplementary Figure 1 | GCCA analysis results detailed on 30 dimensions, suggesting UKB and ADNI group have broader alignment with GCCA 30 dimension subspace than random group**

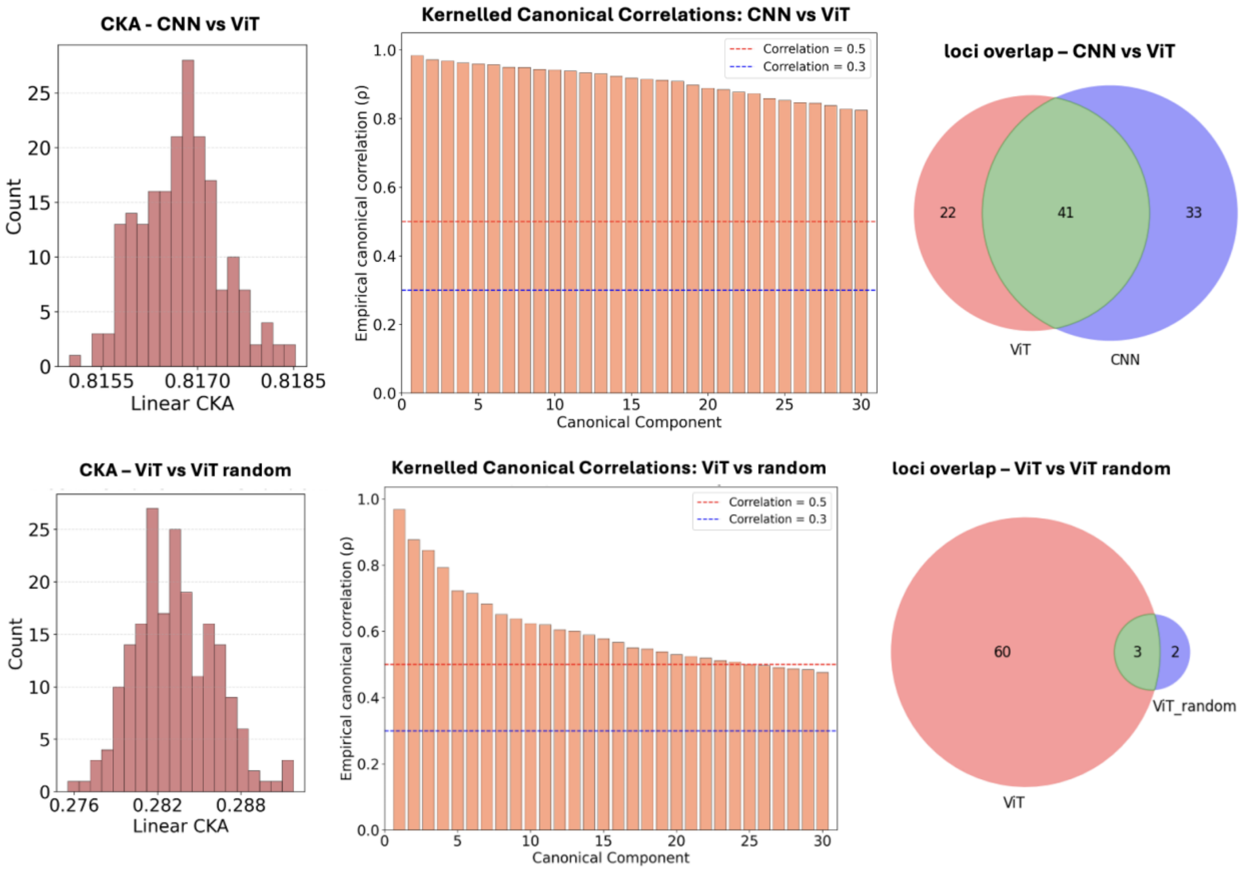


**Supplementary Figure 2 | Model consistency of UDIPs across CNN and ViT.** From left to right: centered kernel alignment (CKA; mean 0.82), kernel canonical correlation analysis (CCA; first 30 canonical correlations; mean 0.91), and loci overlap (0.43). For reference, comparisons between a randomly initialized network and a trained ViT (evaluated on UKB embeddings) yield a mean CKA of 0.28, a mean CCA of 0.61, and loci overlap ratio of 0.05.

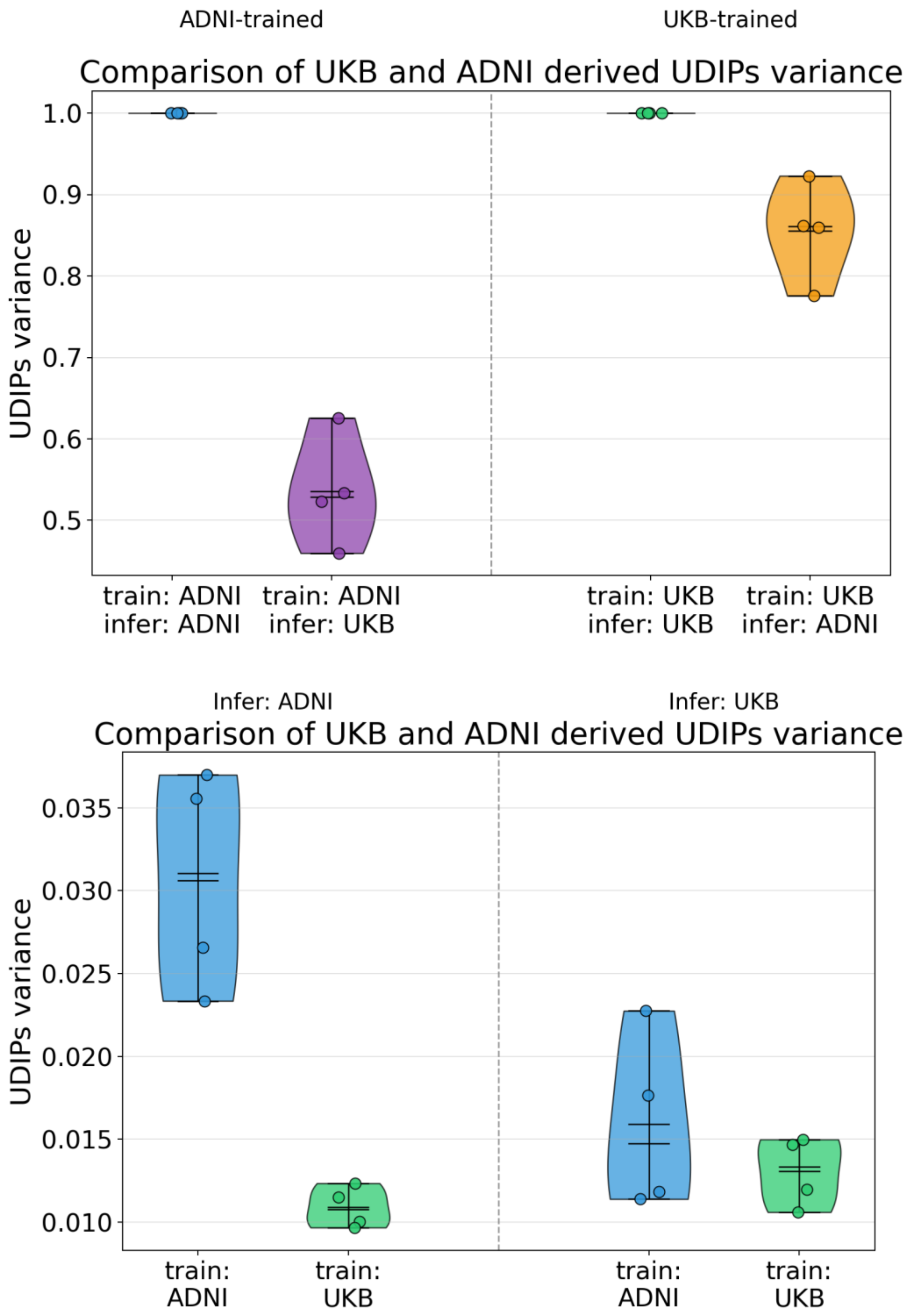


**Supplementary Figure 3 | Top: Training-wise standardized UDIP variance: within- and cross-dataset inference (UKB vs ADNI); Bottom: Inference fixed UDIP variance: train with UKB or ADNI**
